# Regulatory T cells modulate bone marrow stromal cell osteogenesis by activating ROCK-myosin axis and cell contractility

**DOI:** 10.64898/2026.01.16.699899

**Authors:** Eylem Baysal, Azzurra Margiotta, Shuntaro Yamada, Jessica Furriol, Samih Mohamed-Ahmed, Nora Marek, Niyaz Al-Sharabi, Daniela E. Costea, Meadhbh Brennan, Salwa Suliman

## Abstract

Bone regeneration depends not only on the intrinsic signaling in bone marrow-derived mesenchymal stromal cells (BMSC), but also on immune cell-derived cues. Regulatory T cells (Treg) are emerging as immune regulators of tissue repair, yet their direct impact on BMSC osteogenesis remains unexplored. Here, we showed that direct human BMSC-Treg co-culture induced more pronounced osteogenic gene expression and cytokine modulation in BMSC than indirect co-culture. This contact-dependent interaction temporally enhanced BMSC osteogenic gene expression, alkaline phosphatase activity and matrix mineralization by interacting with ROCK-myosin signaling pathway. Exploratory proteomic profiling further revealed enrichment of cytoskeletal, mechanosensitive and osteogenic regulators in BMSC after direct Treg co-culture. Pharmacological inhibition of cell contractility impaired osteogenesis yet Treg co-culture partially restored cytoskeletal integrity and differentiation. These findings identify novel mechanisms behind Treg as direct modulators of BMSC mechanobiology and osteogenesis, highlighting their therapeutic potential in bone regeneration.

## 1. Introduction

Bone marrow-derived mesenchymal stromal/stem cells (BMSC) play a pivotal role in bone regeneration due to their capacity for self-renewal, immunomodulation and osteogenic potential (Dalle Carbonare et al., 2025). Their functionalities are tightly regulated by the dynamic interplay of biochemical signals, mechanical stimuli, cytoskeletal remodeling and crosstalk with immune cells (Dalle Carbonare et al., 2025). However, the influence of immune cells, particularly those of the adaptive immune system, on BMSC cytoskeletal organization during osteogenesis remains largely unexplored.

Regulatory T cells (Treg) have emerged as key modulators of tissue regeneration (Gazdic et al., 2018; Lei et al., 2015; Loffredo et al., 2024). Treg migrate to the injury site and modulate local progenitor cells, promoting healing across systems such as muscle, skin and bone (Li et al., 2018). Local delivery of CD4^+^CD25^hi^Foxp3^+^ Treg enhanced musculoskeletal repair in murine models (Nayer et al., 2024). Treg also supported muscle satellite cell activation and differentiation *in vivo* (Castiglioni et al., 2015). Similarly, Treg facilitated hair follicle regeneration in skin via Jagged1-Notch signaling (Ali et al., 2017).

In bone regeneration, Treg have been mostly studied for their capacity to suppress osteoclastogenesis. This immunoregulatory function is mediated through both soluble factors and contact-dependent mechanisms, including cytotoxic T lymphocyte-associated antigen 4 mediated engagement of CD80/CD86 on osteoclast precursors, which induces apoptosis and attenuates osteoclastogenesis (Kim et al., 2007; Zaiss et al., 2007). While this has established a foundational role for Treg in maintaining skeletal homeostasis, it represents only one facet of Treg function in osteogenesis. Emerging evidence suggests that Treg may influence bone formation by regulating the activity of BMSC. *In vivo* studies demonstrated that Treg enhanced fracture healing and supported BMSC-mediated bone formation, largely through the suppression of local pro-inflammatory cytokines (Liu et al., 2011; Liu et al., 2015). In addition, Treg improved bone parameters in osteogenesis imperfecta by reducing activated T cell phenotype *in vivo* (Kang et al., 2022).

Some evidence suggests that BMSC can stabilize Foxp3 expression in Treg via CD39-CD73 signaling and mitochondrial transfer (Do et al., 2021). Conversely, Treg have also been hypothesized to influence BMSC osteogenesis through similar purinergic signaling pathways and contact-dependent mechanisms, but these claims remain speculative (Lei et al., 2015). A single co-culture study reported a modest increase in alkaline phosphatase (ALP) activity in BMSC exposed to induced Treg without elucidating the underlying mechanisms (Croes et al., 2016).

Recently, we reported that Treg-derived secretome contains proteins linked to cytoskeletal dynamics, metabolism, and immunomodulation, and significantly enhances BMSC osteogenesis (Baysal et al., 2025). However, it remains unclear whether these effects are driven solely through paracrine signaling, direct cell-cell contact, or a combination of both. Notably, direct BMSC-Treg interactions may serve not only as a mode of communication, but also as a source of intracellular signaling. Key mediators of such contact-dependent signaling include members of the Rho family of GTPases regulate cytoskeletal organization and cell-cell adhesion through downstream effectors including Rho-associated coiled-coil kinases (ROCK) (Strzelecka-Kiliszek et al., 2017). Increasing evidence indicates that the Rho/ROCK axis plays a critical role in modulating MSC osteogenic differentiation (McBeath et al., 2004; Yamada et al., 2023). However, the potential for Treg-mediated cell-contact interactions to influence these pathways in the context of osteogenesis remains largely unexplored.

Given the growing interest in immunomodulation as a strategy to enhance BMSC-based bone regeneration, a deeper understanding on how Treg directly influence BMSC molecular function is both timely and necessary. Therefore, we hypothesize that Treg enhance BMSC osteogenic differentiation predominantly through contact-dependent mechanisms involving cytoskeletal organization and Rho/ROCK signaling.

## 2. Results

### 2.1. Direct BMSC-Treg co-culture elicits distinct osteogenic and cytokine responses in BMSC compared to indirect co-culture

BMSC and Treg were characterized by typical surface marker expression, and Treg maintained stable Foxp3 expression on day 7 when cultured alone or directly co-cultured with BMSC in osteogenic media (OM) **(Figure S1, 2)**. BMSC osteogenic gene expression showed time-dependent changes in direct or indirect Treg co-cultures over 7 days in OM. At day 4, *SP7* expression significantly decreased in both co-cultures compared with BMSC monoculture, whereas *RUNX2* expression significantly increased only in direct co-culture relative to both indirect co-culture and monoculture **(Figure 1A)**. By day 7, BMSC *ALP* expression was significantly upregulated in direct co-culture compared to indirect and monoculture, while *SPP1* expression significantly increased only in direct co-culture compared with monoculture **(Figure 1B)**. Additionally, ALP staining showed a higher trend in direct co-culture compared with BMSC monoculture and indirect co-culture **(Figure 1C, D)**.

**Figure 1.**
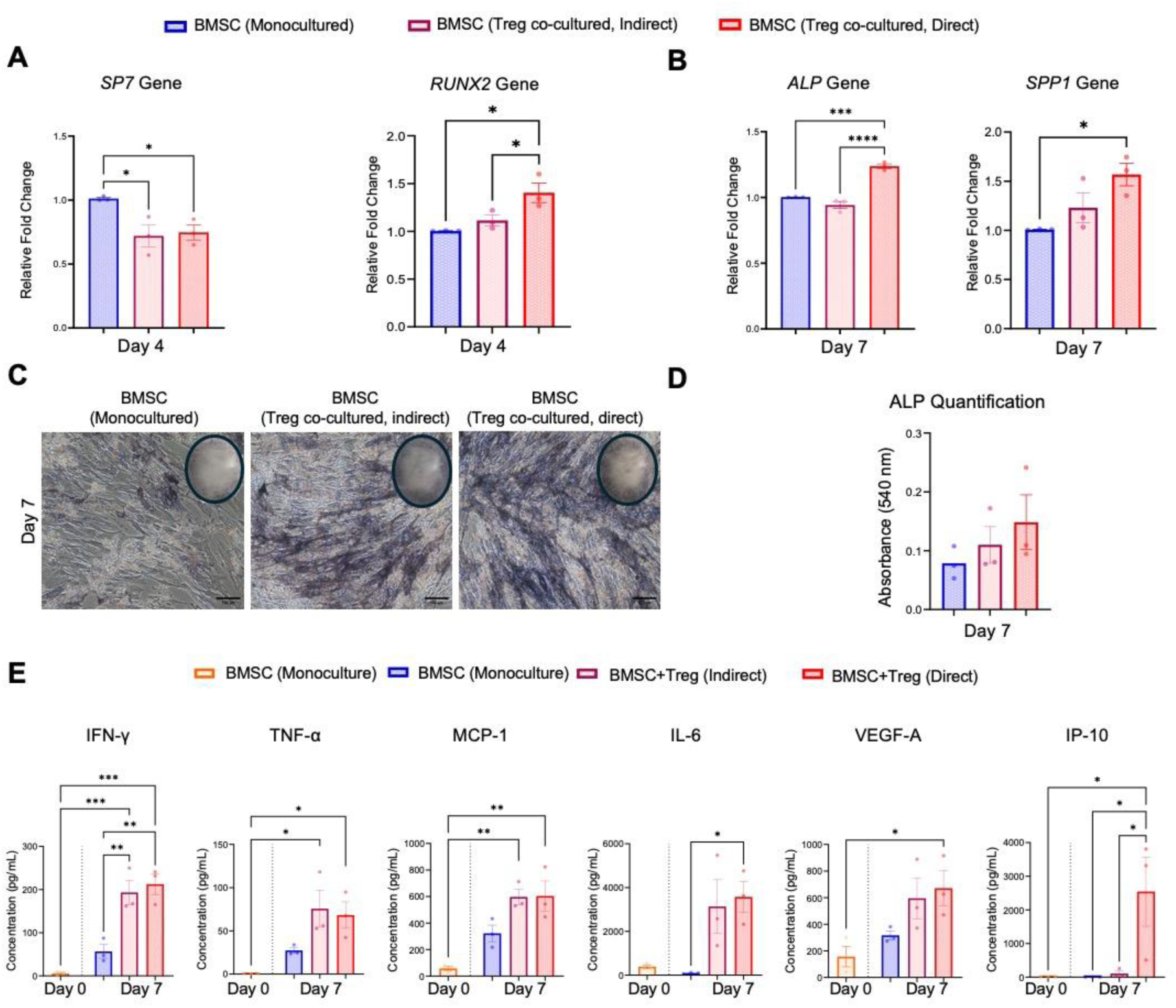
Direct BMSC-Treg co-culture elicits a more prominent osteogenic and immunomodulatory response in BMSC than indirect co-culture. **A.** *SP7* and *RUNX2* expressions as markers of early osteogenic differentiation. **B.** Expressions of *ALP* and *SPP1* as markers of early to mid-stage and mid-to-late stage osteogenic differentiation, respectively. Gene expressions are presented as fold-changes relative to the BMSC monoculture in OM. **C.** Representative image of ALP staining. Scale bar: 100 µm **D.** Quantification of ALP staining. **E.** Cytokine, chemokine and growth factor levels in the culture supernatant from BMSC monoculture, direct and indirect BMSC-Treg co-cultures. Data are presented as mean ± SEM (n = 3). *p ≤ 0.05, **p ≤ 0.01, ***p ≤ 0.001, ****p ≤ 0.0001.

Treg cultured alone in TexMACS or OM for 7 days secreted multiple cytokines, including IL-1Ra, IL-4, IL-5, IL-10 and IL-13, confirming their cytokine-production capacity under both conditions **(Figure S3)**. BMSC with Treg co-culture in OM for 7 days significantly increased IFN-γ levels in both direct and indirect co-culture compared with BMSC monoculture and BMSC at day 0. TNF-α and MCP-1 were significantly upregulated in both co-culture conditions compared to BMSC on day 0. IL-6 levels was significantly higher in direct co-culture compared to BMSC monoculture at day 7, while VEGF-A was significantly increased with Treg co-culture compared to BMSC monoculture at day 0. Notably, IP-10 was significantly increased in direct co-culture only **(Figure 1E)**. The mean absolute concentrations of cytokines from supernatants are summarized in **Table S1.**

Given the more pronounced effect on osteogenic gene expression in BMSC upon direct Treg co-culture compared with indirect (transwell), the direct co-culture model was selected for subsequent investigations.

### 2.2. Direct BMSC-Treg co-culture temporally regulates BMSC osteogenesis under osteogenic conditions

Initially, BMSC exhibited an elongated, spindle-shaped morphology, while Treg showed a spherical shape. In growth media (GM), BMSC retained their typical morphology through day 10. However, in the presence of Treg, BMSC were even more elongated and stretched. Under osteogenic conditions, morphological changes with denser cell layers were evident in BMSC cultures as the cells showed a cuboidal shape. Interestingly, the signs of the matrix deposition occurred as early as day 10 in the co-culture condition, whereas this occurred at day 14 in BMSC cultured alone **(Figure S4A).**

BMSC proliferation occurred earlier with Treg co-culture than in BMSC monoculture in OM. Similar trend was observed in GM **(Figure S4B).** Co-culture with Treg further increased BMSC metabolic activity on day 7 in both conditions, although this did not reach statistical significance **(Figure S4C)**.

Direct Treg co-culture enhanced early BMSC osteogenic differentiation by significantly upregulating *RUNX2* expression on day 1 in GM or OM, and this effect persisted only under OM through day 7 **(Figure 2A)**. In GM, *ALP* levels remained low in both BMSC monoculture, and co-cultures compared to OM, but Treg co-culture induced a higher peak at day 14 than in BMSC monoculture. Similarly, BMSC *ALP* expression peaked earlier and reached the highest level around day 7 when co-cultured with Treg in OM, while ALP gene expression peaked around day 10 in BMSC monoculture in OM **(Figure 2B)**. These transcriptional findings were corroborated by ALP enzymatic activity, which was elevated under OM compared to GM and reached its highest levels in Treg co-cultures in both conditions by day 7 **(Figure 2C)**. Consistent with accelerated differentiation under OM, *BGLAP* expression, a late osteogenic marker, was significantly upregulated in BMSC co-cultured with Treg under osteogenic conditions **(Figure 2D)**.

**Figure 2.**
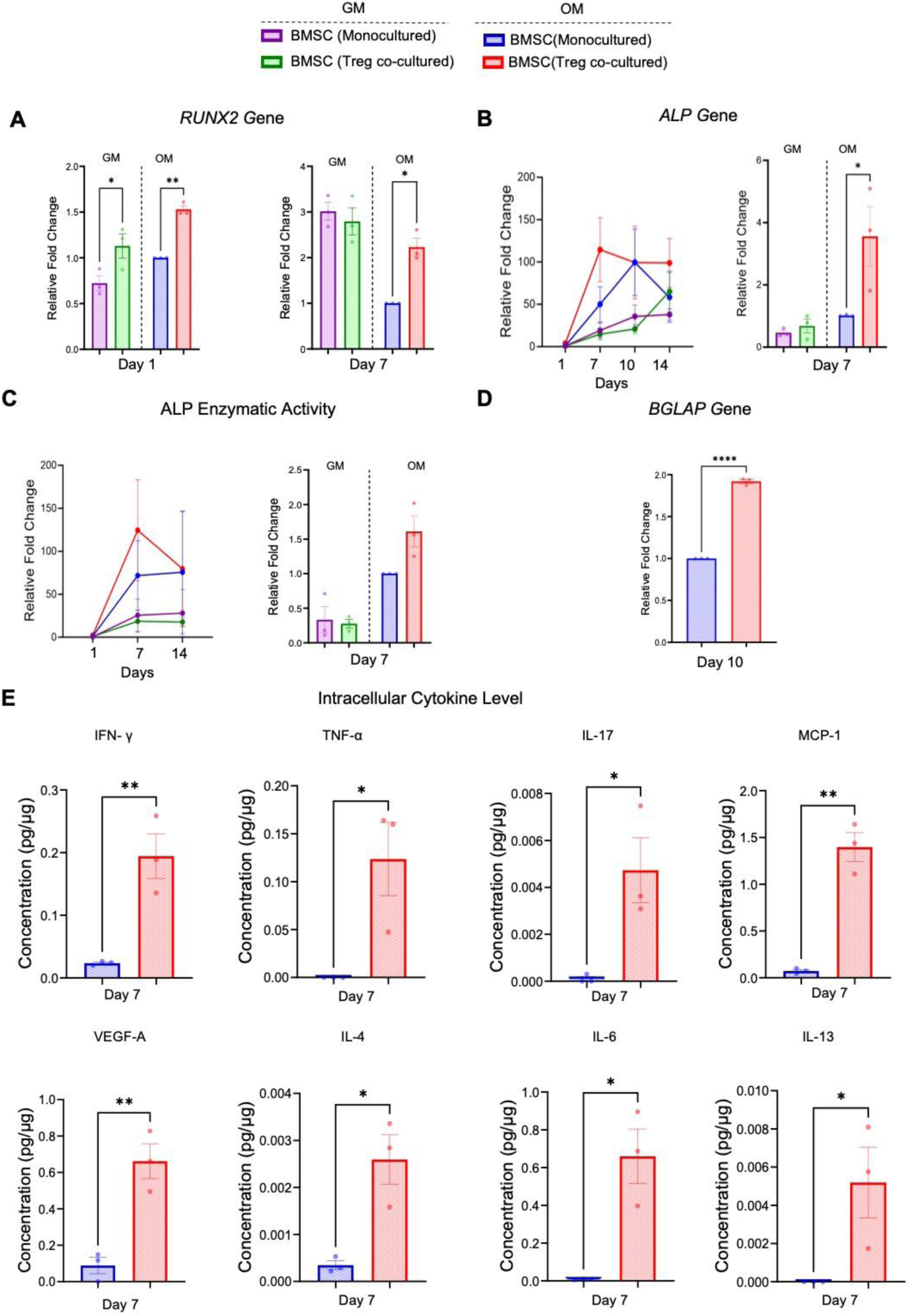
Direct BMSC-Treg co-culture accelerates osteogenic gene expression, ALP activity, and modulates intracellular cytokine production in BMSC. **A.** *RUNX2* expression as early osteogenic differentiation marker. **B.** *ALP* gene expression over time and on day 7. Gene expressions were normalized to BMSC monoculture in GM at day 1 (undifferentiated baseline) or to BMSC cultured in OM at day 7 (osteogenic baseline), respectively. **C.** ALP activity in BMSC over time and on day 7. Data expressed as a fold change relative to BMSC monoculture in GM on day 1 or in OM on day 7, respectively. D. *BGLAP* gene expression as early osteogenic differentiation marker. **E.** Cytokine levels in BMSC lysate normalized to total protein concentration on day 7. Data are presented as mean ± SEM (n = 3). *p ≤ 0.05, **p ≤ 0.01, ****p ≤ 0.0001.

Intracellular cytokine levels in BMSC, including IFN-γ, TNF-α, IL-17, MCP-1, VEGF-A, IL-4, IL-6, and IL-13 were also significantly upregulated with Treg co-culture in OM compared to BMSC monoculture **(Figure 2E)**. A summary of cytokine concentrations measured in the BMSC lysate is in **Table S2**.

BMSC ALP staining progressively increased in OM, with markedly higher staining observed by day 7 with Treg co-culture, which further intensified by day 14 **(Figure 3A)**. At day 7, corresponding to peak BMSC ALP activity and osteogenic gene expression, extracellular vesicle (EV) and matrix vesicles (MV) were isolated from BMSC monocultures and Treg co-cultures to assess vesicle-associated osteogenic activity. The presence of lipid bilayer-enclosed vesicles was confirmed by electron microscopy, with sizes ranging from 100 to 200 nm **(Figure 3B-i)**. Immunophenotyping verified the identity of the isolated vesicles as CD9⁺ populations bound to CD63 beads **(Figure 3B-ii)**. ALP enzymatic activity was significantly elevated in MV from BMSC-Treg co-cultures compared with MV from BMSC monocultures. Within co-cultures, MV exhibited higher ALP activity than EV **(Figure 3B-iii**).

**Figure 3.**
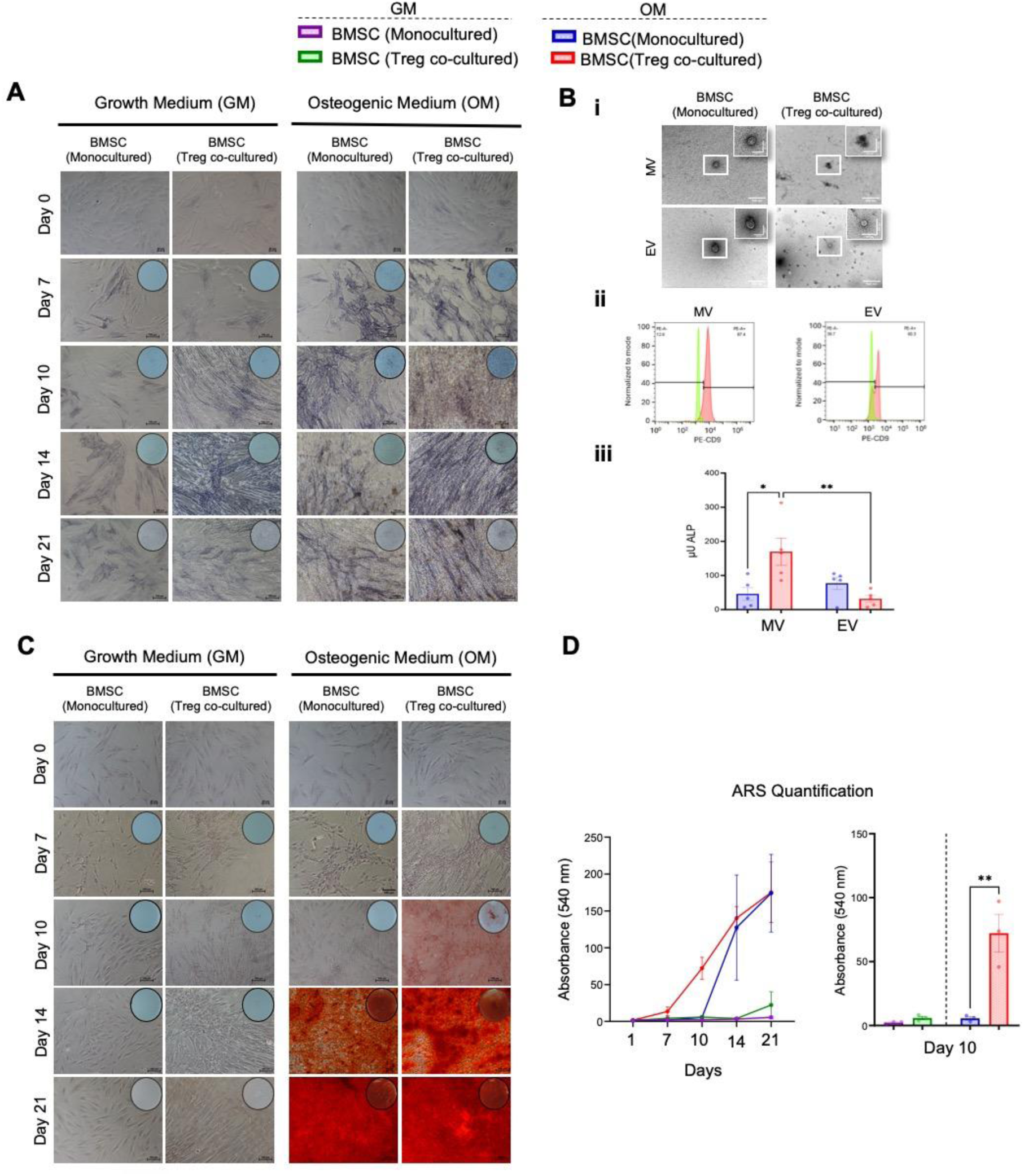
Direct BMSC-Treg co-culture enhances ALP activity and matrix mineralization in BMSC. **A.** Representative ALP staining. Scale bar: 50 µm and 100 µm **B.** Vesicle-associated ALP activity. **(i)** Transmission electron microscopy images of MV and EV from BMSC cultured in OM with or without Treg co-culture for 7 days. Magnification 25.0k×; scale bar: 500 nm. **(ii)** Identification of MV and EV from BMSC cultured in OM as CD9⁺ vesicles isolated using CD63-coated beads and stained with anti-CD9 antibody. **(iii)** ALP enzymatic activity in MV and EV. **C.** Representative Alizarin red S staining. Scale bar: 50 µm and 100 µm **D**. Quantification of the Alizarin red S staining over time and on day 10. Values are expressed as percentages of absorbance. Data are presented as mean ± SEM (n = 3). *p ≤ 0.05, **p ≤ 0.01, ***p ≤ 0.001, ****p ≤ 0.0001.

Early Treg-mediated osteogenic effects were corroborated by enhanced mineralization in BMSC. In OM, mineralized nodules were detectable by day 10 in BMSC-Treg co-cultures, whereas mineralization was only evident by day 14 in BMSC monoculture **(Figure 3C)**. On day 10, BMSC mineralization was significantly increased in BMSC-Treg co-cultures in OM compared with BMSC monocultures **(Figure 3D)**.

Since Treg-mediated BMSC osteogenic differentiation was pronounced at early stages under osteogenic conditions, subsequent analyses were performed under OM to identify the underlying molecular changes.

### 2.3. Direct BMSC-Treg co-culture modulates ROCK-myosin axis in BMSC during osteogenic differentiation

After exclusion of proteins detected in Treg, 145 proteins were exclusively identified as highly upregulated in BMSC with direct Treg co-culture compared to BMSC monoculture **(Figure 4A)**. Functional enrichment analysis of these proteins revealed three major biological clusters **(Figure 4B)**. Cluster 1 was associated with cytoskeletal organization and enriched for processes such as actin cytoskeleton organization, actin filament-based process. Cluster 2 was enriched for osteogenic and tissue developmental processes, including osteoblast differentiation, bone development, and biomineral tissue development. Cluster 3 comprised biological processes related to mechanosensitive and migratory behavior, including cellular response to mechanical stimulus, regulation of cell adhesion, and regulation of cell migration **(Figure 4C)**.

**Figure 4.**
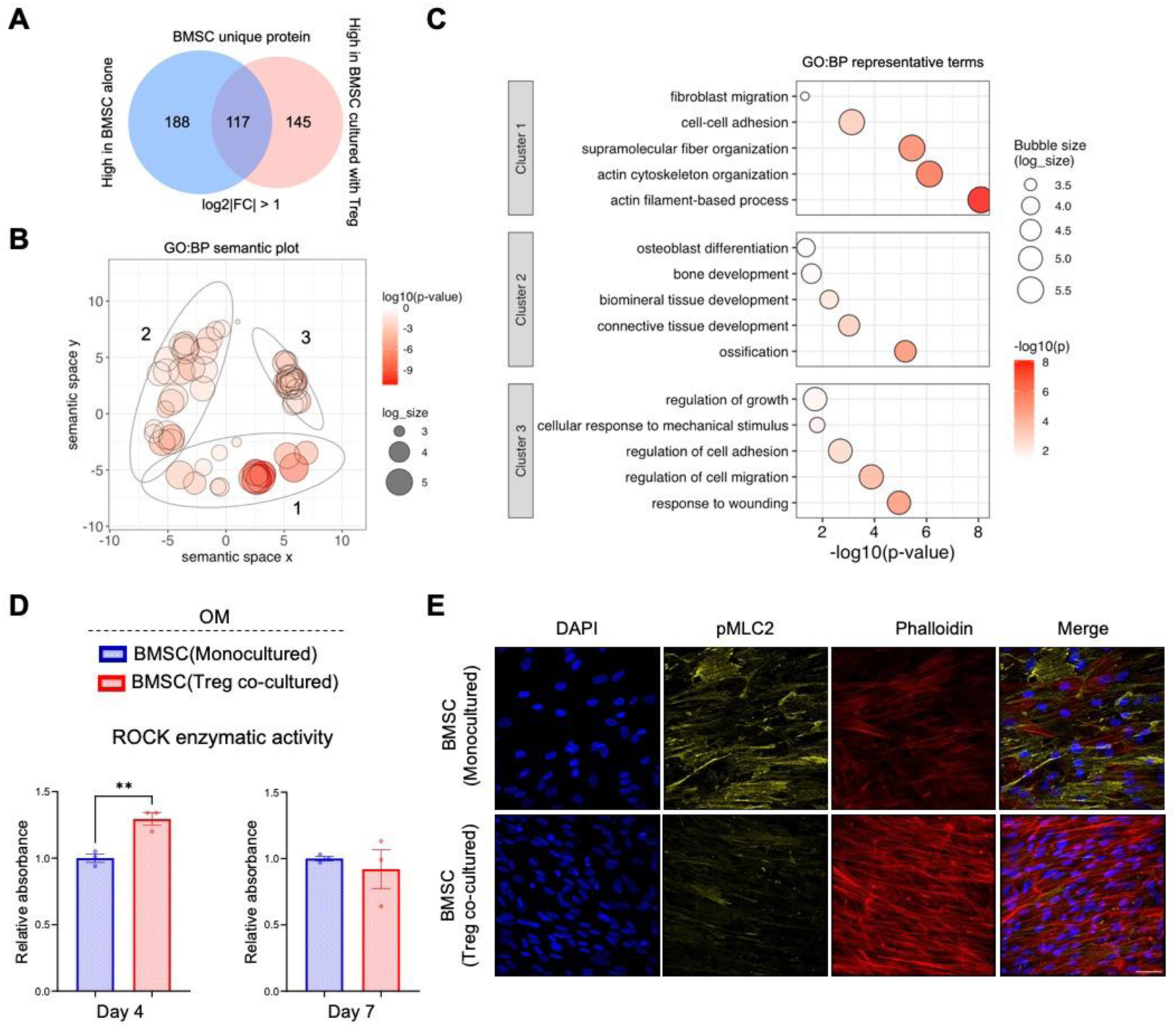
Direct BMSC-Treg co-culture modulates actomyosin signaling and cytoskeletal organization in BMSC. **A.** Venn diagram showing the number of proteins detected exclusively in BMSC and highly upregulated in BMSC after 7 days of direct Treg co-culture compared with BMSC monoculture. **B.** Gene Ontology Biological Process (GO:BP) semantic space plot of enriched terms derived from BMSC-specific proteins that were highly upregulated in BMSC after direct Treg co-culture compared with BMSC monoculture. **C.** Bubble plots of representative GO:BP terms for the three major functional clusters identified. Bubble size reflects the number of proteins associated with each term, and color indicates statistical significance. **D.** ROCK enzymatic activity was expressed as fold change relative to BMSC monoculture in OM. ROCK activity data are presented as mean ± SEM (n = 3). **E.** Representative images showing pMLC2 staining. Images show phalloidin staining for F-actin (red), DAPI staining for nuclei (blue), and p-MLC2 expression (yellow). Scale bar: 50 µm. Data are presented as ± SEM or median (n=3). **p ≤ 0.01

Next, cytoskeletal organization and actomyosin signaling were analyzed in BMSC following direct Treg co-culture. ROCK activity significantly increased on day 4 in BMSC with Treg coculture compared to BMSC monoculture **(Figure 4D)**. At the early stage of co-culture (day 1), BMSC showed increased phosphorylated myosin light chain 2 (pMLC2) and actin intensity under osteogenic conditions compared with BMSC monoculture **(Figure S5)**. By day 7, pMLC2 levels were reduced in BMSC. Notably, BMSC co-cultured with Treg showed a more elongated morphology with a uniformly aligned actin organization compared with BMSC monoculture **(Figure 4E).**

### 2.4. BMSC-Treg direct co-culture enhanced BMSC osteogenesis through ROCK-Myosin II signaling

BMSC monocultures showed dispersed pMLC2 with disorganized, thin actin filaments on day 7 after ROCK (Y-27632) and myosin II (blebbistatin) inhibition (**Figure S6A)**. Under control conditions, BMSC in direct Treg co-culture displayed elongated nuclei and thick, organized actin stress fibers, accompanied by reduced pMLC2 intensity compared with BMSC monocultures on day 7 **(Figure 5A)**. Upon ROCK inhibition in BMSC-Treg co-cultures, Y-27632 increased pMLC2 intensity while decreasing phalloidin intensity. In contrast, blebbistatin treatment further reduced pMLC2 intensity without affecting phalloidin intensity relative to co-cultured without inhibitors **(Figure 5A)**.

**Figure 5.**
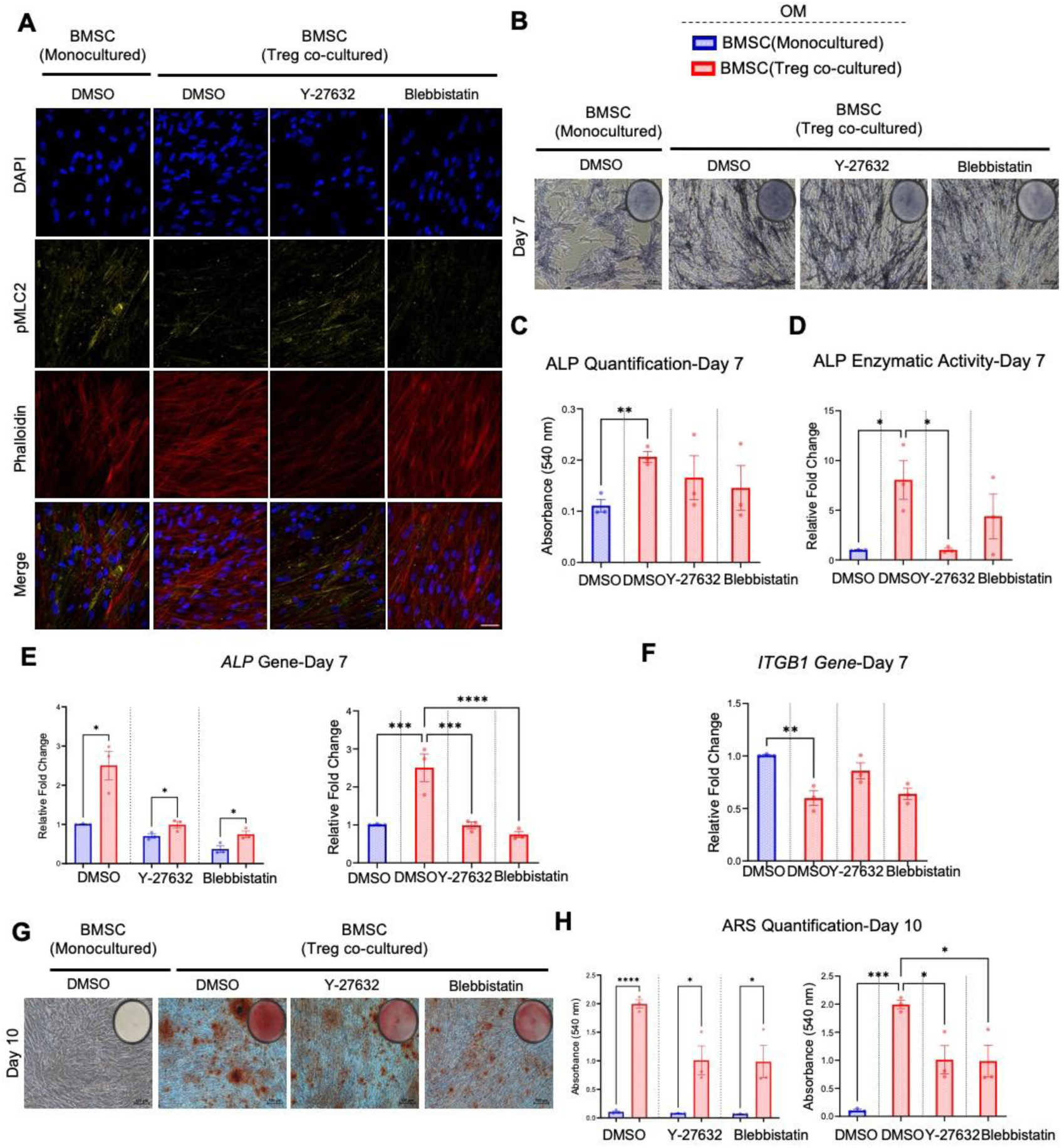
Direct BMSC-Treg co-culture modulated BMSC osteogenic differentiation via ROCK-myosin axis. **A.** Representative images showing DAPI staining for nuclei (blue), p-MLC2 expression (yellow) and F-actin (red) on day 7. Scale bar: 50 µm. **B.** Representative images of ALP staining. Scale bar: 100 µm. **C.** Quantification of ALP staining intensity in BMSC shown in panel **B**. **D.** ALP enzymatic activity in BMSC, expressed as a fold change relative to BMSC monocultured with DMSO in OM. **E.** Expression of *ALP* in BMSC. **F.** Expression of *ITGB1* gene in BMSC. Gene expressions presented as fold-change relative to BMSC monocultured with DMSO in OM. **G.** Representative images of Alizarin red S staining in BMSC. Scale bar: 100 µm. **H.** Quantification of Alizarin Red S staining in BMSC shown in panel **G.** Data are presented as mean ± SEM (n = 3). *p ≤ 0.05, **p ≤ 0.01, ***p ≤ 0.001, ****p ≤ 0.0001.

During osteogenic differentiation, both Y-27632 and blebbistatin significantly reduced ALP staining, quantified enzymatic activity and gene expression in BMSC monocultures on day 7 **(Figure S6B, C, D, E)**. While direct Treg co-culture significantly enhanced these parameters in BMSC compared with BMSC monocultures, the effects were attenuated by both inhibitors **(Figure 5B, C, D, E).** Notably, BMSC ALP activity was significantly decreased by Y-27632, while *ALP* expression was significantly downregulated by both inhibitors in Treg co-cultures compared with Treg co-cultures without inhibition **(Figure 5D, E)**. *ITGB1* expression in BMSC was significantly reduced by Treg co-culture compared with monoculture on day 7 **(Figure 5F)**. However, no significant differences were observed with Y-27632 or blebbistatin, neither in monocultures nor in Treg co-cultures, compared with their respective controls **(Figure 5F; Figure S6F)**.

By day 10, Alizarin Red staining revealed no detectable mineralization in any group cultured without Treg **(Figure S6G)**. On the other hand, Treg co-culture significantly enhanced BMSC mineralization compared to monocultures, but the extent of mineralization was significantly reduced by Y-27632 and blebbistatin treatment **(Figure 5G, H)**.

## 3. Discussion

Treg have recently gained attention not only for their immunosuppressive roles but also for their emerging capacity to promote tissue regeneration through direct interactions with tissue-resident progenitor cells (Lei et al., 2015). In the context of bone regeneration, we provide a comprehensive mechanistic insight into how Treg influence the osteogenic differentiation of BMSC.

We previously showed that Treg secretome enhances BMSC osteogenesis and comprises a diverse repertoire of cytokines (Baysal et al., 2025). Many of these factors including IL-10, VEGF-A, IFN-γ, TNF-α are known to promote osteoblastogenesis or modulate the milieu to favor regeneration (Zhao et al., 2022). Our current findings support a dual mechanism by which Treg modulate BMSC osteogenesis through both soluble mediators and direct cell-cell contact. Although the cellular source of cytokines detected in the supernatant cannot be distinguished, both direct and indirect Treg co-culture altered the BMSC cytokine environment, suggesting the creation of an osteoinductive niche. For instance, IFN-γ, TNF-α and MCP-1 levels were elevated in the supernatant from both co-culture conditions. It has been reported previously that IFN-γ knockdown impaired *RUNX2* expression and osteogenesis (Duque et al., 2009). Also, TNF-α at low concentrations enhanced osteogenic markers, including *ALP*, *SPP1*, and *RUNX2*, in murine MSC (Huang et al., 2011). The role of MCP-1 in osteogenesis has also been shown *in vivo*, as MCP-1⁻/⁻ mice failed to upregulate key osteogenic genes in response to parathyroid hormone stimulation (Siddiqui et al., 2021).

Indeed, soluble factors initiated osteogenic signaling in both co-cultures, and direct BMSC-Treg contact provided additional, distinct cues. VEGF-A and IL-6 levels were specifically elevated in direct co-culture, both are known to stimulate osteoblast proliferation and matrix formation (Huang et al., 2010; Xie et al., 2018). Interestingly IP-10 (CXCL10) was uniquely upregulated in direct co-culture. Although its role in osteogenesis remains unexplored, CXCL10 is known to recruit CXCR3⁺ Treg (Li et al., 2020), and enhance MSC migration (Hermans et al., 2024), suggesting a potential feedback loop that sustains BMSC-Treg interactions and amplifies regenerative signaling.

Beyond secreted factors, intracellular cytokine accumulation in BMSC lysates, particularly IL-17A, IL-4, and IL-13, suggest active synthesis or uptake of osteoimmune mediators. IL-13 and IL-17A have been shown to induce *RUNX2* expression, ALP activity, and promote mineralization in mouse mesenchymal precursors (Khalmuratova et al., 2019), aligning with enhanced *RUNX2* and *ALP* expression in direct co-culture observed in our study. Together, these findings indicate that while soluble factors from Treg initiate osteogenic signaling in BMSC, only direct co-culture robustly activated osteogenic gene expression.

Indeed, direct co-culture promoted and sustained *RUNX2* expression in BMSC, particularly under OM, highlighting the synergistic effect of osteogenic supplements and Treg derived signals. *In vivo*, bone regeneration is governed by a complex interplay of cellular and molecular cues that drive osteogenesis after injury (Dalle Carbonare et al., 2025). Our *in vitro* model mimics this environment by combining biochemical induction with immunomodulatory support from Treg. Consistent with early lineage commitment, Treg co-cultures promoted an ALP activation and subsequent mineral deposition. This progression underscores the capacity of Treg to drive matrix mineralization, not merely initiate differentiation.

Matrix mineralization is a tightly regulated process, beginning with the deposition of amorphous calcium phosphate, which is converted into hydroxyapatite, the mineral component of bone (Lotsari et al., 2018; Sugimoto et al., 2023). This transition is initiated within matrix vesicles released by osteoblasts into the collagen-rich extracellular matrix (Ansari et al., 2021). ALP, localized on these vesicles, plays a pivotal role in the calcification processes (Golub, 2009). Higher ALP activity in MV after Treg co-culture suggests Treg modulate BMSC vesicle composition, thereby accelerating mineralization as early as day 10.

The biogenesis of MV is closely linked to cytoskeletal remodeling (Bottini et al., 2018), particularly via the ROCK signaling (Strzelecka-Kiliszek et al., 2017), which regulates vesicle release, cell shape and matrix interactions. The enrichment of proteins which is associated with cytoskeletal organization and mechanosensitive signaling in BMSC following Treg co-culture points to involvement of ROCK-dependent pathways in Treg-mediated osteogenesis. Mechanistically, ROCK enhance actomyosin contractility by phosphorylating MLC (Strzelecka-Kiliszek et al., 2017). The increased pMLC2 together with enhanced F actin organization observed as early as day 1, followed by elevated ROCK activity by day 4 in BMSC after Treg co-culture, further supports that Treg-derived cues robustly engage this ROCK signaling pathway, thereby promoting vesicle formation, cytoskeletal remodeling and accelerated mineral deposition. On the other hand, reduced pMLC2 expression with denser and organized actin fibers was observed in BMSC after Treg co-culture by day 7, suggesting that a temporal shift from contractile activity toward cytoskeletal stabilization and matrix organization. Such cytoskeletal arrangement is known to influence cell mechanical properties, including stiffness, which in turn provides instructive cues for MSC lineage differentiation (Khan et al., 2020). MSC respond to mechanical stiffness by forming prominent actin stress fibers, and increasing cytoskeletal tension, which enhance *RUNX2* and *BGLAP* expressions to accelerate mineralization (Shih et al., 2011). The enrichment of proteins associated with osteogenic differentiation, together with the enhanced osteogenic gene expression and mineralization in BMSC after Treg co-culture support that Treg-derived signals mimic stiffness cues to promote osteogenesis via ROCK-mediated cytoskeletal reorganization.

As anticipated, both ROCK and myosin II inhibition impaired actomyosin-contractility and reduced osteogenic progression, consistent with previous studies (Kilian et al., 2010; Yamada et al., 2023). Although Treg-mediated BMSC osteogenesis was attenuated under inhibitory condition, Treg co-culture still partially preserved *ALP* activity, and mineral deposition. These findings indicate Treg promote BMSC osteogenesis through both contractility-dependent cytoskeletal remodeling and complementary contractility-independent mechanisms.

Nevertheless, certain aspects should be considered in this study. The use of primary cells from multiple donors strengthens biological relevance, this also introduces inter-donor variability. Additionally, the co-culture system integrates soluble factors, EV and direct cell-cell contact, reflecting physiological complexity; however, it does not fully mimic the broader immune microenvironment present *in vivo*, thereby limiting precise mechanistic dissection, and direct clinical translation. While pharmacological inhibition provides valuable mechanistic insights, these approaches are inherently reversible and less specific. Therefore, future studies using genetic knockdown strategies and systematically dissecting the individual contributions of soluble mediators and vesicular cargo will be instrumental to define the key signals to orchestrate BMSC osteogenic differentiation.

Altogether, our findings reveal a multifaceted role for Treg in promoting osteogenic differentiation of BMSC through a regulation of cytoskeletal architecture. These cytoskeletal changes provide instructive cues for osteogenic commitment, and Treg-derived signals in direct co-culture robustly enhanced osteogenic gene expression and matrix mineralization in BMSC, partially mediated by ROCK-myosin signaling and actomyosin contractility. These findings provide novel mechanistic insights into how Treg regulate mesenchymal stromal cell behavior and highlight their potential as therapeutic modulators for bone regeneration.

## 5. Material and Methods

### 5.1. Ethical approval and cell sources

BMSC were isolated from donors undergoing standard surgical procedures at Haukeland University Hospital, following informed consent and ethical approval from Regional Committees for Medical Research Ethics in Norway (Regional Ethical Committee reference number: 2020/7199/REK sør-øst C). Peripheral blood collection for Treg isolation was approved by Haukeland Blood Bank Bergen, Norway and no ethical approval was required.

### 5.2. Characterization and multilineage differentiation of BMSC

BMSC were isolated from the iliac crest bone marrow aspirates, as previously described (Mohamed-Ahmed et al., 2018), expanded in Dulbecco’s Modified Eagle’s medium (DMEM) (Invitrogen, USA) supplemented with 10% (v/v) fetal bovine serum (FBS) (GE Healthcare, USA) and 1% (v/v) penicillin/streptomycin (P/S) (GE Healthcare) and characterized as negative for CD34, HLA-DR, and CD45, and positive for CD73, CD105, and CD90 using a BD Accuri C6 Flow cytometer (BD Biosciences) before performing further experiments **(Figure S1A)** .

BMSC multilineage differentiation was also confirmed by osteogenic and adipogenic differentiation, as previously described (Mohamed-Ahmed et al., 2018). Osteogenic differentiation was evaluated on day 7 by ALP staining, and on days 14 and 21 by Alizarin Red S staining **(Figure S1B)**. Adipogenic differentiation was assessed on days 14 by Oil Red O staining **(Figure S1C)**.

### 5.3. Expansion and characterization of regulatory T cells

Treg were isolated from human peripheral blood using the MACSxpress Whole Blood Treg Isolation Kit and expanded with CD3/CD28 MACSiBead Expansion Kit (Miltenyi Biotec, Germany), following the manufacturer’s instructions. Treg were cultured in expansion medium including TexMACS medium (Miltenyi Biotec) supplemented with 5% AB serum (Sigma-Aldrich), and 500 U/mL IL-2 (Miltenyi Biotec). After 9 days, Treg were phenotyped using the Human Treg Detection Kit (Miltenyi Biotec), analyzed on a ID7000™ Spectral Cell Analyzer (Spectral Cell Analyzer, USA) and characterized as CD45^⁺^CD4^⁺^CD25^⁺^CD127^dim/-^Foxp3^⁺^ **(Figure S2A)**.

### 5.4. Comparing direct and indirect co-culture of BMSC and Treg

BMSC were cultured alone or co-cultured with Treg either directly in the same well or indirectly using a Transwell system (0.4 µm pore size, Sarstedt, Germany) at a 1:3 ratio for 7 days in OM (DMEM supplemented with 5% AB serum, 1% P/S, 173 µM L-ascorbic acid 2-phosphate, 10 nM dexamethasone, and 10 mM β-glycerophosphate) for an initial comparison. Culture media were freshly prepared, supplemented with 50 U/mL IL-2 (Miltenyi Biotec) at each medium change, and refreshed twice per week. All reagents were from Sigma-Aldrich, USA, unless otherwise stated.

Before all downstream analyses of BMSC, Treg were removed from co-cultures by washing three to five times with Dulbecco’s Phosphate-Buffered Saline (DPBS; Thermo Fisher Scientific, USA) supplemented with 0.5 mM EDTA (Sigma-Aldrich).

### 5.5. Temporal analysis of BMSC under direct co-culture with Treg

For subsequent in-depth analyses, BMSC and Treg were co-cultured directly for up to 21 days, either in GM (DMEM with 5% AB serum and 1% P/S) or OM with 50 U/mL IL-2. Morphological alterations were observed over 21 days using an inverted light microscope (Nikon Eclipse TS100, Japan). Different time-course experiments were performed under GM and OM to assess BMSC proliferation, metabolic activity, osteogenic gene expressions, ALP activity and extracellular matrix mineralization. Furthermore, mechanistic studies, including vesicle ALP activity, cytokine profiling from lysates, total proteome profiling and functional analysis of the Rho/ROCK-Myosin II pathway were performed under OM.

### 5.6. BMSC proliferation and metabolic activity

BMSC proliferation was evaluated under GM or OM, either alone or co-cultured with Treg directly, by using the PicoGreen dsDNA assay, according to the manufacturer’s instructions on days 1, 7, 10, 14, and 21. BMSC metabolic activity was assessed after 7 days using the Alamar Blue Cell Viability Reagent (Thermo Fisher Scientific), according to the manufacturer’s instructions.

### 5.7. BMSC gene expression

Total RNA from BMSC was extracted using RNeasy Kit (Qiagen, Germany) and 100-300 ng was reverse-transcribed into cDNA using the High-Capacity cDNA Reverse Transcription Kit (Thermo Fisher Scientific), following the manufacturer’s instructions. Expressions of *GAPDH*, *RUNX2*, *SP7*, *ALP*, *SPP1*, *BGALP* and *ITGB1* were evaluated at specific time points using quantitative real-time polymerase chain reaction with TaqMan Fast Universal PCR Master Mix (2×) (Thermo Fisher Scientific) and analyzed using the comparative C(T) method **(Table S3)**. Data were normalized to *GAPDH* expression.

### 5.8. BMSC ALP activity

For ALP staining, BMSC were fixed with 4% paraformaldehyde (Merck, USA), incubated with BCIP/NBT Liquid Substrate System (Sigma-Aldrich) for 15 min and imaged by an inverted light microscope (Nikon Eclipse TS100). The dye was dissolved in 100 mM cetylpyridinium chloride (Sigma-Aldrich) overnight and absorbance was measured at 540 nm with a Varioskan LUX Multimode Microplate Reader (Thermo Fisher Scientific).

For ALP enzymatic activity, BMSC were lysed in 0.1% Triton X-100 (Sigma-Aldrich)/DPBS following two freeze-thaw cycles, sonicated and mixed 1:1 with the Alkaline Phosphatase Yellow (pNPP) Liquid Substrate System for ELISA (Sigma-Aldrich) at 37°C for 30 min. Absorbance was measured at 405 nm. Cell number was quantified using the Quant-iT PicoGreen dsDNA assay (Thermo Fisher Scientific), following the manufacturer’s instructions. ALP activity was normalized to cell number.

### 5.9 BMSC-derived vesicle ALP activity

#### 5.9.1 Extracellular and matrix vesicle isolation

BMSC were cultured in OM alone or co-cultured directly with Treg. On day 7, after 24h of incubation in serum-free media, the supernatant or cell layers were collected separately, as previously described (Skelton et al., 2023). Supernatant was centrifuged at 600 × g for 5 min and collected for EV isolation. The cell layer was collected using a scraper after trypsinization, centrifuged at 3000 × g for 5 min and supernatant was collected for MV isolation. Supernatants were concentrated using Amicon Ultra-15 filters (100 kDa MWCO, Merck), centrifuged at 3000 × g for 15 min at 4 °C, then EV and MV were isolated using size exclusion chromatography with a qEVoriginal 35 nm column (Izon Science, New Zealand). Fractions 1-6 (3 mL, non-vesicular proteins) were discarded, and fractions 7-11 (2.5 mL, vesicle-enriched) were pooled and stored at -80 °C.

#### 5.9.2. Characterization of EV and MV

The morphology of EV and MV were assessed using transmission electron microscopy (TEM). Vesicle suspensions were fixed with 2% glutaraldehyde (1:1) for 30 min, then applied to a formvar carbon-coated copper grid (200 mesh) for 10 min, stained with 1.5% uranyl acetate for 12 s and imaged by TEM (Hitachi High-Technologies, Japan) (Rana et al., 2022).

The EV and MV were immunophenotyped using anti-CD63-coated Dynabeads (Invitrogen). Vesicles were incubated with beads and resuspended with 1% bovine serum albumin (BSA; Sigma-Aldrich) in 0.2 µm filtered DPBS overnight at 4 °C with rotation, then stained with anti-human CD9 antibody or IgG1 isotype control (BD Biosciences) for 1 h. After washing, samples were resuspended in 0.1% BSA/DPBS and analyzed using the BD Accuri6 flow cytometer.

ALP activity of vesicles was measured using the same pNPP-based enzymatic assay described in Section 5.8. Samples were incubated with substrate solution (1:1) at 37 °C for 30 min and absorbance was measured at 405 nm, expressed as relative to a standard curve of calf intestinal alkaline phosphatase (Sigma-Aldrich).

### 5.10. BMSC extracellular mineralization

BMSC were fixed with 4% paraformaldehyde for 15 min and incubated with 2% Alizarin Red S (pH 4.2; Sigma-Aldrich) for 45 min, then washed and dried overnight. Images were visualized using an inverted light microscope. For quantification, samples were incubated with 100 mM cetylpyridinium chloride for 30 min, and absorbance was read at 544 nm with microplate reader.

### 5.11. Cytokine profiling from supernatants or BMSC lysates

Cytokine concentrations were measured in supernatants collected from BMSC at day 0 or at day 7 in OM (monocultured or co-cultured with Treg directly or indirectly), as well as from Treg cultured alone in expansion medium or OM for 7 days. After Treg removal from direct co-cultures by washing with DPBS-EDTA, BMSC were lysed using RIPA buffer with protease and phosphatase inhibitors (Thermo Fisher Scientific). Cytokine levels were then measured and normalized to protein concentration determined by the BCA Protein Assay Kit (Thermo Fisher Scientific).

A multiplex fluorescent bead-based immunoassay was performed using the Bio-Plex Pro Human Cytokine 27-plex Assay (Bio-Rad, USA) on the Luminex platform (Luminex, USA), following the manufacturer’s instructions.

### 5.12 Total proteome analysis

#### 5.12.1 Sample preparations

Proteomic analyses were performed on day 7 using BMSC from a single donor cultured alone in OM, BMSC directly co-cultured with Treg in OM, Treg cultured alone in expansion medium. Cells were lysed in RIPA buffer with protease and phosphatase inhibitors. Protein samples were processed using the SP3 (single-pot, solid-phase–enhanced sample preparation) method. Samples were adjusted to equal volumes and reduced with 10 mM dithiothreitol for 20 min at 60 °C, followed by alkylation with 8 mM iodoacetamide for 1 h at room temperature in the dark. Proteins were bound to a mixture of hydrophobic and hydrophilic Sera-Mag SpeedBeads at a bead-to-protein ratio of 10:1 (w/w) by addition of ethanol to a final concentration of 70%, followed by incubation for 7 min at 24 °C with shaking. Bead-bound proteins were washed three times with 80% ethanol and digested on-bead in 50 µL of 100 mM ammonium bicarbonate containing 1 mM CaCl₂ and trypsin (0.2 µg/µL) for 16 h at 37 °C.

After digestion, samples were centrifuged, placed on a magnetic rack, and the supernatant was transferred to a new tube. Beads were then resuspended in 50 µL of 0.5 M NaCl, sonicated, centrifuged again, and placed on a magnetic rack. The resulting supernatant was combined with the initial digest supernatant (final volume ∼100 µL). Peptides were subsequently desalted using 2 mg OASIS C18 96-well plates and analyzed using an Orbitrap Eclipse Tribrid mass spectrometer in data independent acquisition.

#### 5.12.2 Proteome Data Analysis

Raw mass spectrometric data processing was performed using Proteome Discover Software (Thermo-Fisher). Raw mass spectrometry data were searched against the human UniProtKB/Swiss-Prot database (release 2022_05, downloaded from https://www.uniprot.org). Data were transferred to Perseus version 1.5.5.3 for normalization and further analysis.

#### 5.12.3 Bioinformatics analysis

To minimize the risk of cross-contamination between co-cultured cell populations, proteins detected in Treg were excluded from analysis and only BMSC-associated proteins were retained for exploratory downstream bioinformatic analysis. A Venn diagram was generated in R to show the overlap and uniqueness of protein sets. Gene Ontology Biological Process (GO:BP) enrichment analysis was performed using g:Profiler (term size < 1000) on proteins exclusively upregulated (log_2_FC > 1) in BMSC co-cultured with Treg (n = 145) compared with BMSC alone. Enriched GO:BP terms were subsequently reduced by semantic similarity using REVIGO and visualized as a two-dimensional semantic space plot clustered by K-means clustering on R. Representative GO:BP terms from each cluster were further displayed as bubble plots.

### 5.13. Assessment of ROCK-Myosin II pathway

BMSC were cultured in OM with or without Treg under direct co-culture conditions. Cytoskeletal imaging was performed on days 1 and 7, and ROCK enzymatic activity was measured on days 4 and 7.

For inhibition experiments, BMSC were cultured in OM supplemented with either 10 µM Y-27632 (ROCK inhibitor; Biotechne, UK) or 10 µM Blebbistatin (myosin II inhibitor; Sigma-Aldrich). Dimethyl sulfoxide (DMSO; Sigma-Aldrich) was used as vehicle control, as previously described (Yamada et al., 2023). In parallel, Treg were cultured alone or co-cultured with BMSC directly in OM with DMSO, Y-27632, or Blebbistatin for 7 days, and characterized using flow cytometry. All media were supplemented with 50 U/mL IL-2 and inhibitors and refreshed every 2 days.

#### 5.13.1 Cytoskeletal imaging

BMSC were fixed in 4% paraformaldehyde, permeabilized in 0.1% Triton X-100/DPBS for 15 min, and blocked in DPBS containing 10% normal goat serum (Abcam, UK)/0.1% Tween-20 (Sigma-Aldrich) for 1 h. Cells were incubated overnight at 4°C with anti-phospho-MYL9 (Ser19) (1:250, Invitrogen) in blocking buffer to detect pMLC2. The following day, samples were incubated for 1 h with CoraLite Plus 647-Goat Anti-Mouse Recombinant Secondary Antibody (1:500, Proteintech, USA), together with DAPI (1:1000, Thermo Fisher Scientific) and Phalloidin-Atto 565 (1:500, Sigma-Aldrich). Images were acquired using a Dragonfly 505 confocal spinning disc microscope (Andor Technologies, Northern Ireland) with identical microscope and detector settings for all experimental conditions.

#### 5.13.2 ROCK enzymatic assay

ROCK activity was measured using the ROCK Activity Assay Kit (Merck), following the manufacturer’s instructions. BMSC were lysed in mammalian cell lysis buffer with protease and phosphatase inhibitor (Abcam) and added to the MYPT1-procoated wells. The kinase reaction was initiated by the supplied mixture for 1 h, followed by incubation with anti-phospho-MYPT1 (Thr696) antibody for 1 h and an HRP-conjugated secondary antibody for another1 h. Absorbance was measured at 450 nm using a microplate reader.

### 5.14. Statistical Analysis

BMSC from three different independent donors were used at passages 3-5 after their characterization. For each donor, experiments were performed with three to five technical replicates per condition. Data are expressed as mean ± standard error of the mean (SEM) from three independent donors (n=3), unless otherwise specified in the figure legend. Statistical analyses were performed using GraphPad Prism 10.0.2 software (GraphPad, USA). For comparisons between BMSC co-cultured directly with Treg and BMSC alone, a Student’s t-test was used. For multiple comparisons, one-way ANOVA, followed by Tukey’s post hoc test was performed. P values ≤ 0.05 were considered statistically significant.

## Resource availability

### Lead contact

Request for further information, resources and data analysis should be directed to lead contact Salwa Suliman

### Material availability

This study did not generate unique reagents.

### Data and code availability

All data are provided in figures, tables and supplementary information or are otherwise available upon request to the lead contact e-mail.

## Acknowledgements

Flow cytometry was conducted at the Flow & Mass Cytometry Core Facility, Department of Clinical Science, University of Bergen. Proteomics was performed at the Proteomics Unit (PROBE), Department of Biomedicine, University of Bergen. Imaging was carried out at the Molecular Imaging Center (MIC), Department of Biomedicine, University of Bergen. The authors acknowledge Niyaz Al-Sharabi for discussions during proteomics analysis.

## Author Contributions

**EB** and **AM**: Methodology, Investigation, Data curation, Formal Analysis, Visualization, Writing – original draft. **SY**: Methodology, Investigation, Formal Analysis, Visualization,

Writing – review & editing. **JF**: Methodology, Formal Analysis, Writing – review & editing. **SMH**: Resources, Writing – review & editing. **NM**: Investigation, Writing – review & editing. **NAS:** Investigation, Writing – review & editing. **DEC**: Methodology, Writing – review & editing **MB**: Methodology, Formal Analysis, Writing – review & editing **SS**: Conceptualization, Methodology, Supervision, Funding Acquisition, Writing – review & editing. All authors reviewed the final version of the manuscript.

