## Supplemental Information for "Regulatory T cells modulate bone marrow stromal cell osteogenesis by activating ROCK-myosin axis and cell contractility"

1. Center of Translational Oral Research (TOR) - Tissue Engineering Group, Department of Clinical Dentistry, Faculty of Medicine, University of Bergen, Bergen, Norway
2. Centre for Craniofacial & Regenerative Biology, Faculty of Dentistry, Oral & Craniofacial Sciences, King's College London, London, UK
3. Department of Medicine, Haukeland University Hospital, Bergen, Norway.
4. The Gade Laboratory for Pathology and Center for Cancer Biomarkers (CCBIO) Department of Clinical Medicine, University of Bergen, Bergen, Norway.
5. Department of Pathology, Haukeland University Hospital, Bergen, Norway.
6. Biomedical Engineering, School of Engineering; Discipline of Advanced Therapeutics, School of Medicine; CÚRAM, the Research Ireland Centre for Medical Devices, University of Galway, Ireland.
7. These authors contributed equally
8. Lead contact

* Corresponding author

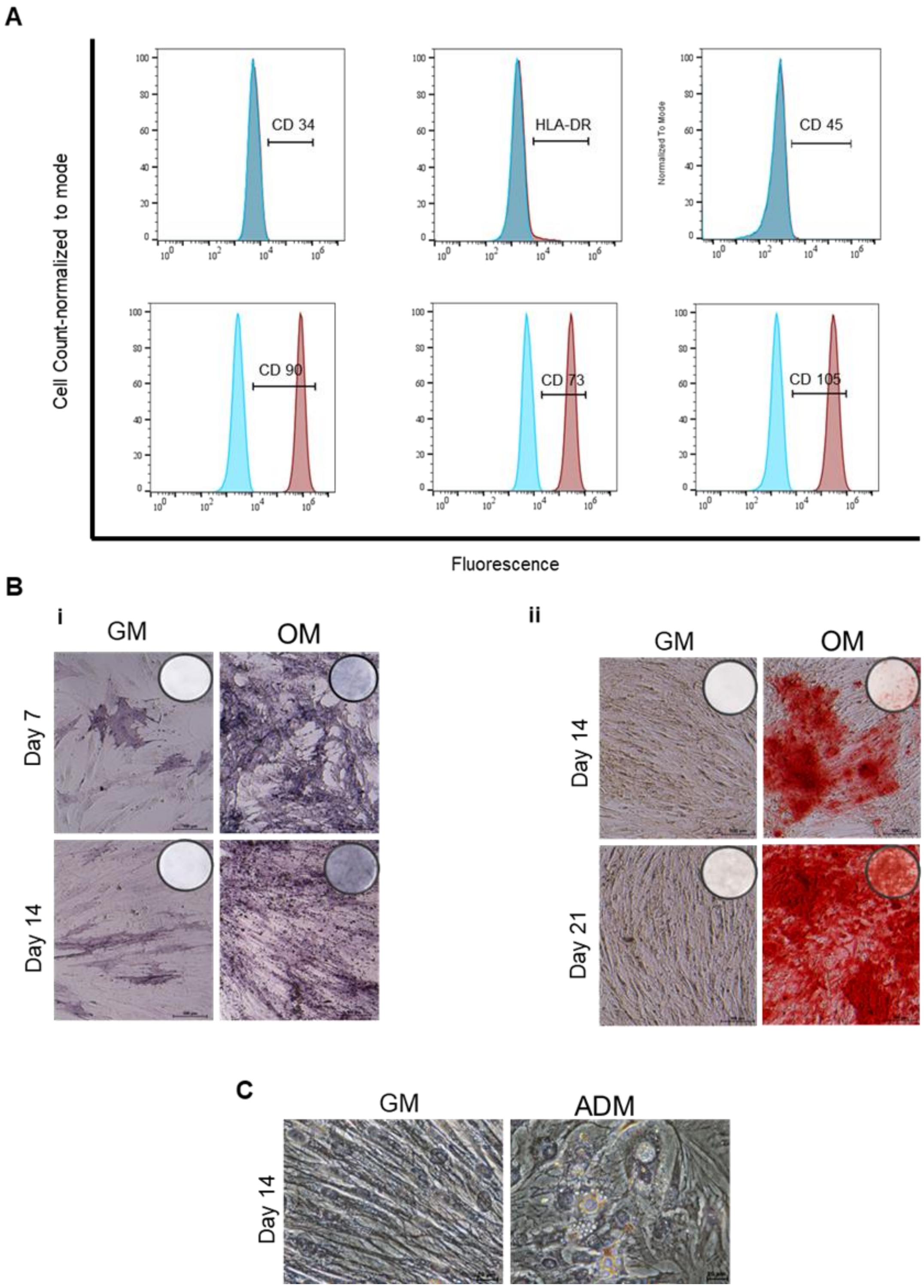


**Figure S1. Characterization and multilineage differentiation of BMSC. A**.BMSC were characterized as negative for CD34, HLA-DR, CD45 and positive for CD90, CD73, CD105 to confirm their stemness phenotype. **B.** Osteogenic differentiation of BMSC. (i) ALP staining on days 7, 14, (ii) Extracellular matrix mineralization by Alizarin red S staining on days 14 and 21. Magnifications 10× scale bars 100 µm. **C.** Adipogenic differentiation of BMSC. Lipid droplets were stained with Oil Red O. Magnifications 20× scale bars 20 µm.

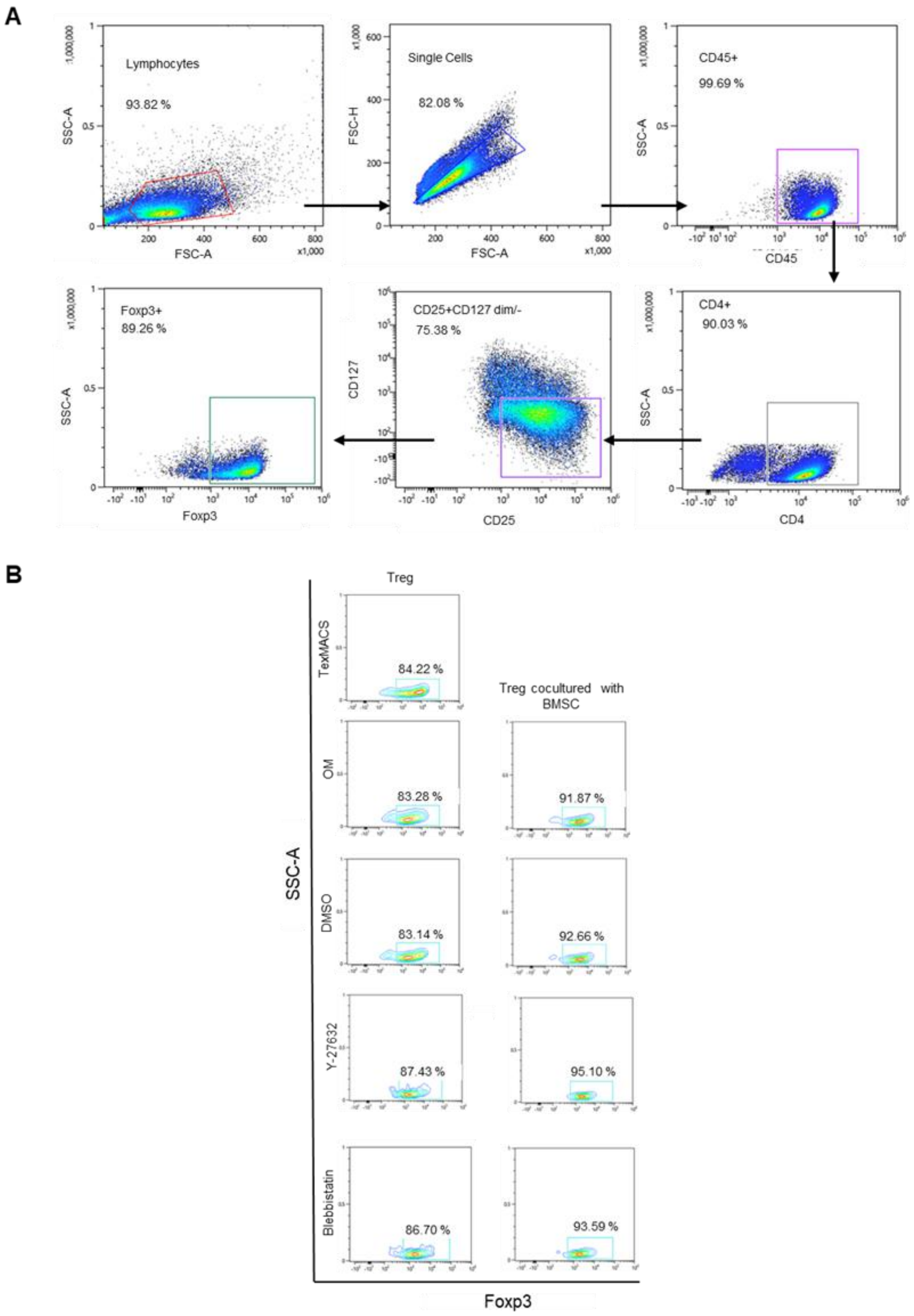


**Figure S2. Phenotypic and functional characterization of Treg. A**. Treg were characterized as CD45$^+$CD4$^+$CD127dim/–CD25$^+$Foxp3$^+$. **B.** Foxp3 expression in Treg cultured alone in TexMACS or OM and after direct co-culture with BMSC treated with DMSO (vehicle control), Y-27632 (ROCK inhibitor), or Blebbistatin (myosin II inhibitor) under OM for 7 days.

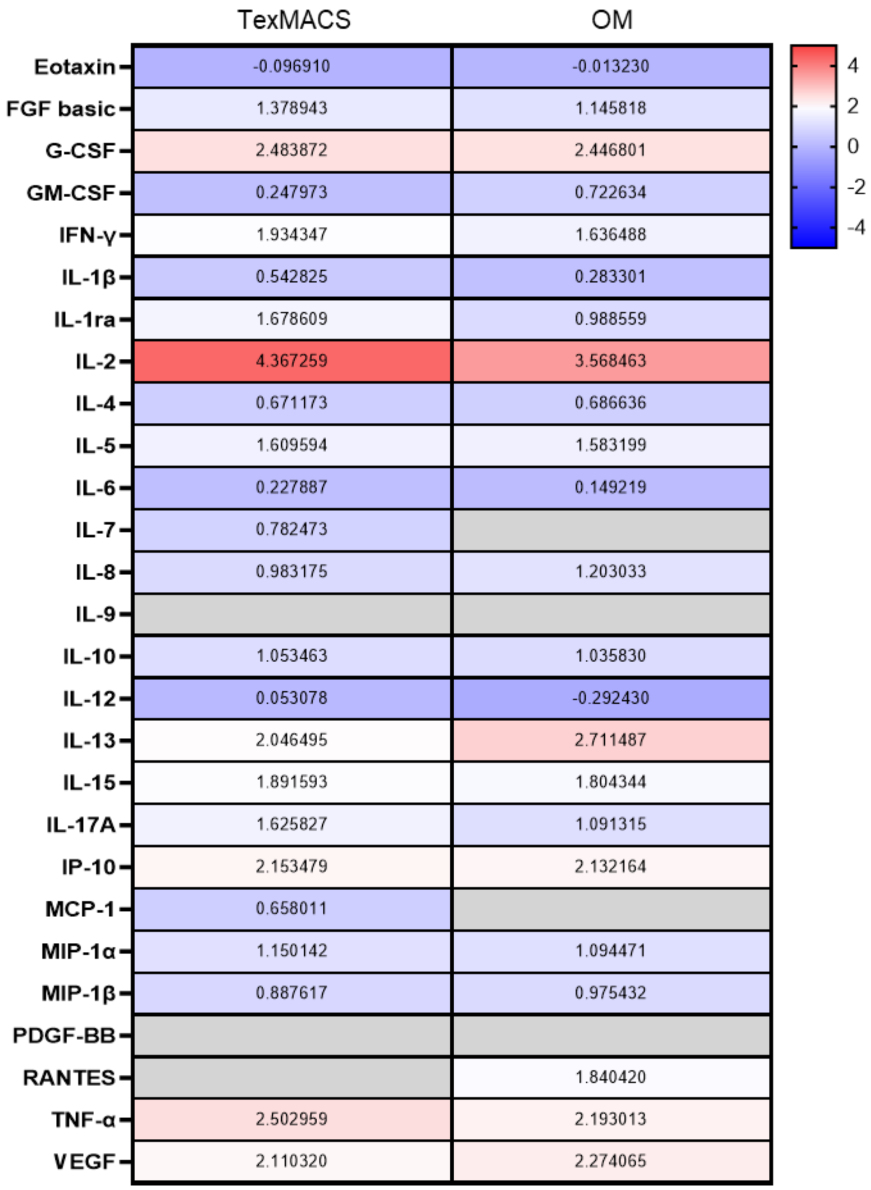


**Figure S3. Cytokine profile of Treg cultured alone in TexMACS or OM for 7 days.** Cytokine concentrations were measured in pg/mL and are shown as $\log_{10}$-transformed values in a heat map. The color scale ranges from –4 (blue, low expression) to +4 (red, high expression).

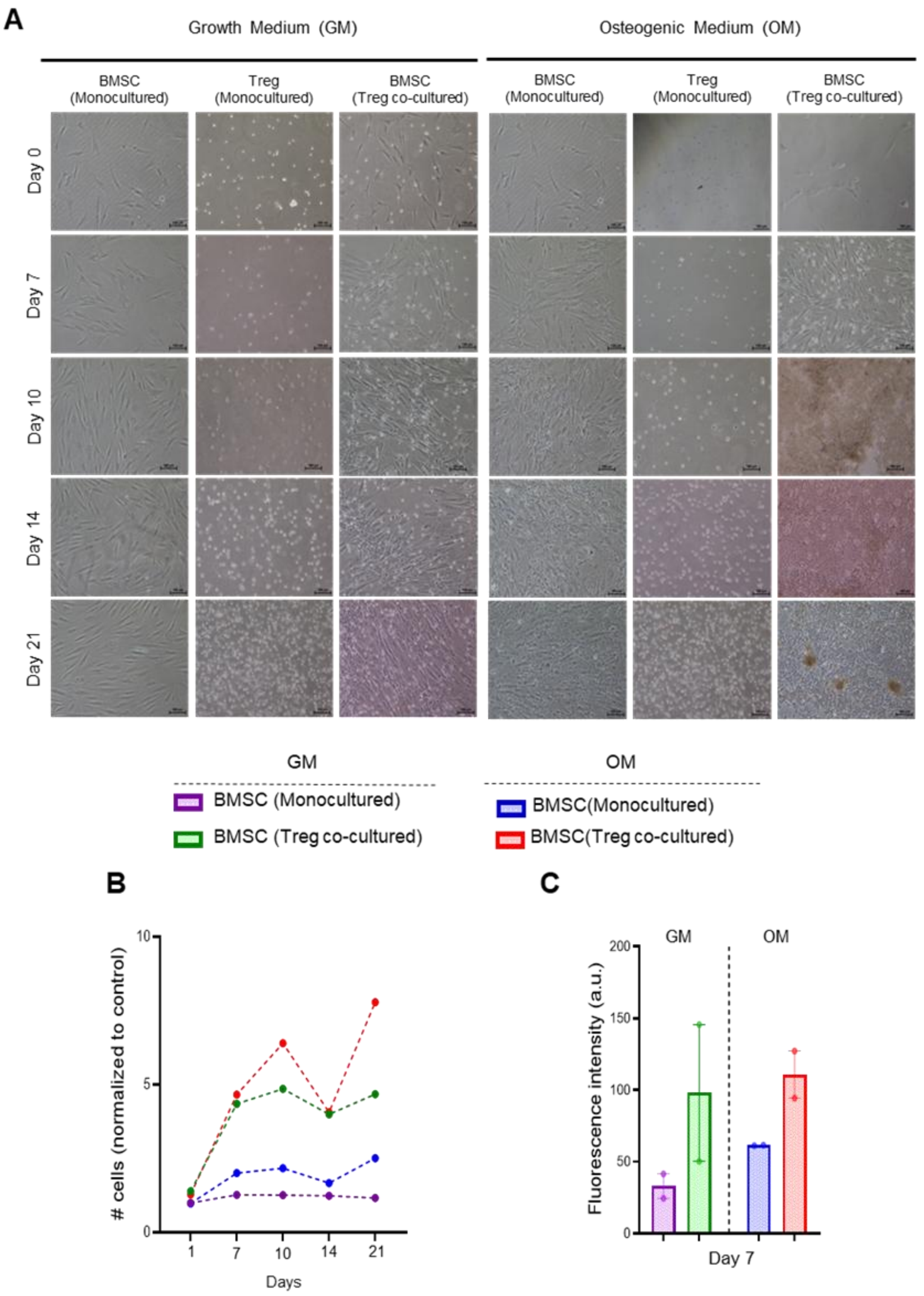


**Figure S4. Morphological assessment of BMSC in the presence of Treg over time**. **A**. Phase-contrast images of BMSC and Treg cultured alone or directly co-cultured under GM and OM over 21 days. Magnification 10×; scale bar: 100 μm. **B**. Proliferation of BMSC cultured alone or with Tregs in GM and OM over 21 days. Data are from three independent donors and presented as fold change relative to BMSCs cultured in GM at day 1. **C**. Metabolic activity of BMSC cultured alone or with Treg in GM and OM on day 7. Data are presented as mean ± SEM. (n = 2).

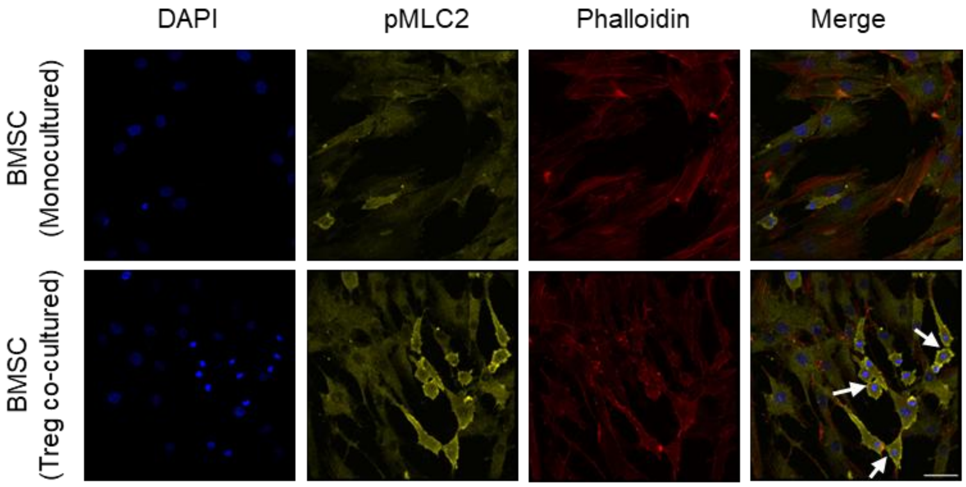


**Figure S5.** Representative images showing phalloidin staining for F-actin (red), DAPI staining for nuclei (blue), and p-MLC2 expression (yellow). White arrows show cells in division and cytokinesis. Scale bar: 50 μm.

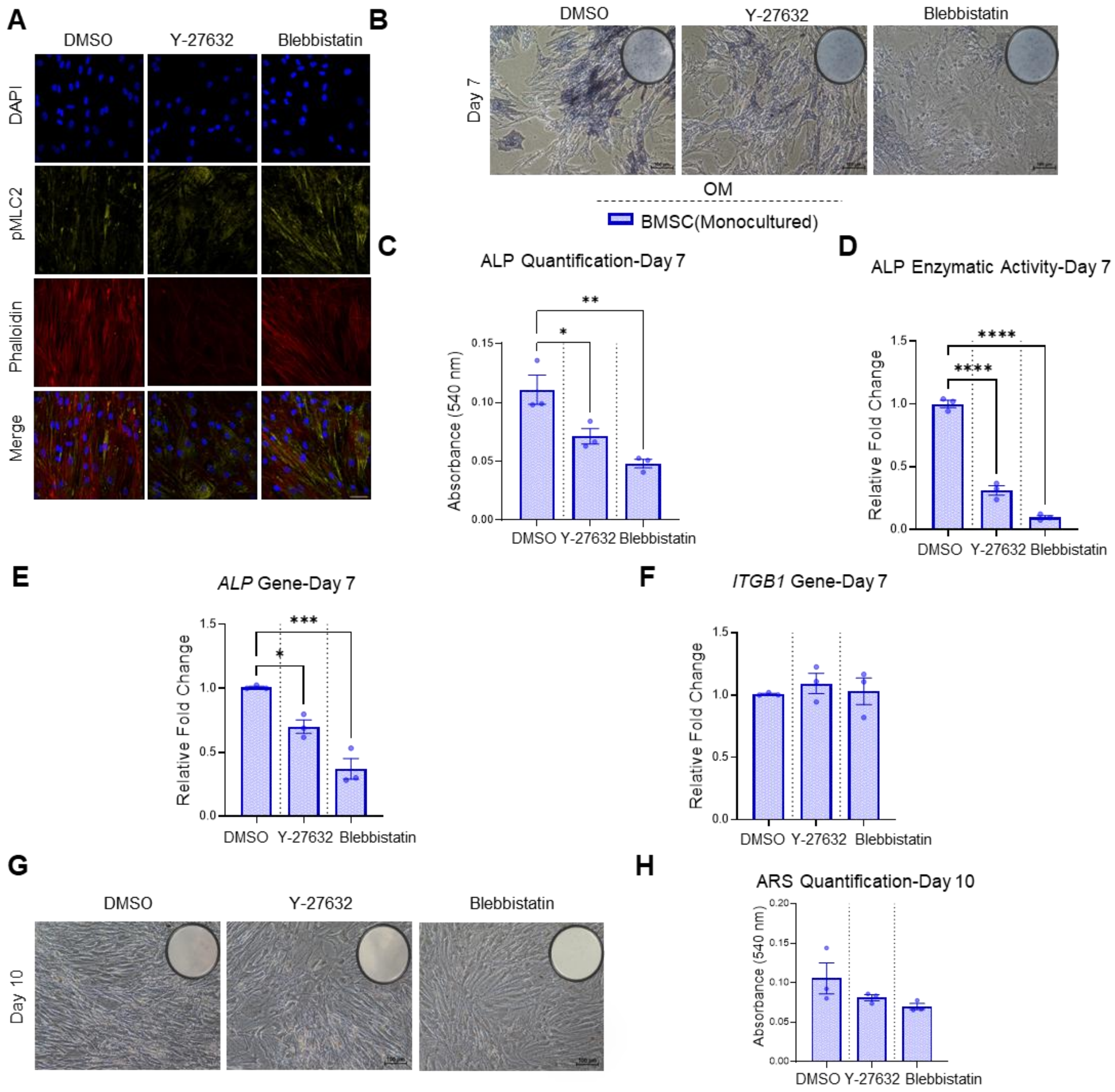


**Figure S6. Impact of ROCK and myosin II inhibition on osteogenic differentiation of BMSC-Treg co-culture under OM. A.** Representative images showing pMLC2 staining. Images show phalloidin staining for F-actin (red), DAPI staining for nuclei (blue), and p-MLC2 expression (yellow). Scale bar: 50 μm. **B.** Representative images of ALP staining. Scale bar: 100 μm. **C.** Quantification of ALP staining intensity in BMSC shown in panel B. **D.** ALP enzymatic activity in BMSC, expressed as a fold change relative to cells cultured in DMSO. **E.** Expression of *ALP* gene in BMSC. **F.** Expression of *ITGB1* gene in BMSC. **G.** Representative images of Alizarin Red S staining in BMSC. Scale bar: 100 μm. **H.** Quantification of Alizarin Red S staining in BMSC shown in panel F. Gene expressions presented as fold-change relative to BMSC-treated with DMSO in OM. Data are presented as mean ± SEM. (n = 3). *$p \leq 0.05$, **$p \leq 0.01$, ***$p \leq 0.001$, ***$p \leq 0.0001$.

**Table S1.** Absolute concentrations of analytes from supernatants of BMSC under different co-culture conditions for 7 days.

| **Chemokine** | | | | |
|---|---|---|---|---|
| Marker | Group | Average Level (pg/ml) | SEM | P-value |
| IL-8 | BMSC (Day 0) | 34.32 | 10.65 | 0.02 |
| | BMSC | 8.977 | 1.424 | |
| | BMSC+Treg (Transwell) | 17441 | 1.041 | |
| | BMSC+Treg (Direct) | 31.99 | 3.737 | |
| IP-10 | BMSC (Day 0) | 0 | 0 | 0.02 |
| | BMSC | 0 | 0 | |
| | BMSC+Treg (Transwell) | 106 | 78.19 | |
| | BMSC+Treg (Direct) | 2542 | 1028 | |
| Eotaxin | BMSC (Day 0) | 0 | 0 | 0.09 |
| | BMSC | 0 | 0 | |
| | BMSC+Treg (Transwell) | 45887 | 8.114 | |
| | BMSC+Treg (Direct) | 34.21 | 19.38 | |
| MCP-1 | BMSC (Day 0) | 58.34 | 15.79 | 0.002 |
| | BMSC | 322.2 | 62.26 | |
| | BMSC+Treg (Transwell) | 596.6 | 58.22 | |
| | BMSC+Treg (Direct) | 604.1 | 112.9 | |
| MIP-1a | BMSC (Day 0) | 0 | 0 | 0.3373 |
| | BMSC | 0.07333 | 0.02404 | |
| | BMSC+Treg (Transwell) | 17.95 | 13.99 | |
| | BMSC+Treg (Direct) | 7.413 | 4.904 | |
| MIP-1b | BMSC (Day 0) | 0 | 0 | 0.5886 |
| | BMSC | 0 | 0 | |
| | BMSC+Treg (Transwell) | 22 | 21.58 | |
| | BMSC+Treg (Direct) | 24.28 | 24.28 | |
| RANTES | BMSC (Day 0) | 0 | 0 | 0.489 |
| | BMSC | 21.31 | 21.27 | |
| | BMSC+Treg (Transwell) | 249.8 | 215.3 | |
| | BMSC+Treg (Direct) | 285.6 | 232.5 | |
| **Growth Factor** | | | | |
| Marker | Group | Average Level (pg/ml) | SEM | P-value |
| bFGF | BMSC (Day 0) | 0 | 0 | 0.0235 |
| | BMSC | 2.177 | 0.4667 | |
| | BMSC+Treg (Transwell) | 16.89 | 6.298 | |
| | BMSC+Treg (Direct) | 15.13 | 3.817 | |
| G-CSF | BMSC (Day 0) | 0 | 0 | 0.1596 |
| | BMSC | 14.92 | 1.938 | |
| | BMSC+Treg (Transwell) | 216.4 | 127 | |
| | BMSC+Treg (Direct) | 106.2 | 39.15 | |
| GM-CSF | BMSC (Day 0) | 0 | 0 | 0.4121 |
| | BMSC | 0 | 0 | |
| | BMSC+Treg (Transwell) | 4.963 | 4.349 | |
| | BMSC+Treg (Direct) | 1.387 | 1.221 | |
| PDGF-bb | BMSC (Day 0) | 0 | 0 | >0.99 |
| | BMSC | 0 | 0 | |
| | BMSC+Treg (Transwell) | 0 | 0 | |
| | BMSC+Treg (Direct) | 0 | 0 | |
| VEGF | BMSC (Day 0) | 156 | 76.24 | 0.0327 |

| | | | | |
|---|---|---|---|---|
| | BMSC | 316.6 | 33 | |
| | BMSC+Treg (Transwell) | 595.1 | 153.4 | |
| | BMSC+Treg (Direct) | 670.8 | 130.4 | |
| **Pro-inflammatory & Anti-inflammatory Marker** | | | | |
| Marker | Group | Average Level (pg/ml) | SEM | P-value |
| IL-1b | BMSC (Day 0) | 0 | 0 | 0.0239 |
| | BMSC | 0.07333 | 0.07333 | |
| | BMSC+Treg (Transwell) | 3.513 | 1.428 | |
| | BMSC+Treg (Direct) | 3.157 | 0.7767 | |
| IL-2 | BMSC (Day 0) | 0 | 0 | 0.0015 |
| | BMSC | 883.9 | 42.01 | |
| | BMSC+Treg (Transwell) | 548.6 | 146.2 | |
| | BMSC+Treg (Direct) | 775.7 | 146.3 | |
| IL-6 | BMSC (Day 0) | 386.1 | 72.24 | 0.0152 |
| | BMSC | 94.8 | 19.37 | |
| | BMSC+Treg (Transwell) | 3135 | 1225 | |
| | BMSC+Treg (Direct) | 3570 | 704.2 | |
| IL-7 | BMSC (Day 0) | 0 | 0 | 0.1316 |
| | BMSC | 0 | 0 | |
| | BMSC+Treg (Transwell) | 4.937 | 2.541 | |
| | BMSC+Treg (Direct) | 3.467 | 1.858 | |
| IL-9 | BMSC (Day 0) | 0 | 0 | >0.99 |
| | BMSC | 0 | 0 | |
| | BMSC+Treg (Transwell) | 0 | 0 | |
| | BMSC+Treg (Direct) | 0 | 0 | |
| IL-12 (p70) | BMSC (Day 0) | 0.06667 | 0.06667 | 0.4547 |
| | BMSC | 0.8367 | 0.08253 | |
| | BMSC+Treg (Transwell) | 1.663 | 0.607 | |
| | BMSC+Treg (Direct) | 32143 | 0.5014 | |
| IL-15 | BMSC (Day 0) | 0 | 0 | 0.0426 |
| | BMSC | 44805 | 7.648 | |
| | BMSC+Treg (Transwell) | 64.94 | 20.70 | |
| | BMSC+Treg (Direct) | 53.17 | 8.324 | |
| IL-17 | BMSC (Day 0) | 0 | 0 | 0.0121 |
| | BMSC | 0.4233 | 0.1267 | |
| | BMSC+Treg (Transwell) | 12663 | 3.402 | |
| | BMSC+Treg (Direct) | 18111 | 1.651 | |
| IFN-ɣ | BMSC (Day 0) | 11810 | 2.744 | 0.0002 |
| | BMSC | 56.68 | 15.77 | |
| | BMSC+Treg (Transwell) | 193.1 | 28.51 | |
| | BMSC+Treg (Direct) | 212.5 | 23.92 | |
| TNF-α | BMSC (Day 0) | 0 | 0 | 0.0112 |
| | BMSC | 27.34 | 3.322 | |
| | BMSC+Treg (Transwell) | 75.66 | 21.28 | |
| | BMSC+Treg (Direct) | 68.27 | 15.19 | |
| IL-1ra | BMSC (Day 0) | 0 | 0 | 0.062 |
| | BMSC | 0 | 0 | |
| | BMSC+Treg (Transwell) | 45745 | 15.17 | |
| | BMSC+Treg (Direct) | 20.47 | 2.135 | |
| IL-4 | BMSC (Day 0) | 0 | 0 | 0.0293 |
| | BMSC | 0.2433 | 0.1955 | |

| | | | | |
|---|---|---|---|---|
| | BMSC+Treg (Transwell) | 4.867 | 1.736 | |
| | BMSC+Treg (Direct) | 5.903 | 2.088 | |
| IL-5 | BMSC (Day 0) | 22890 | 6.756 | 0.0046 |
| | BMSC | 45680 | 1.461 | |
| | BMSC+Treg (Transwell) | 147.3 | 33.7 | |
| | BMSC+Treg (Direct) | 144.4 | 33.03 | |
| IL-10 | BMSC (Day 0) | 0 | 0 | 0.298 |
| | BMSC | 0 | 0 | |
| | BMSC+Treg (Transwell) | 45967 | 4.608 | |
| | BMSC+Treg (Direct) | 2.783 | 1.403 | |
| IL-13 | BMSC (Day 0) | 0 | 0 | 0.0919 |
| | BMSC | 0 | 0 | |
| | BMSC+Treg (Transwell) | 412.7 | 208.1 | |
| | BMSC+Treg (Direct) | 318.5 | 130.4 | |

**Table S2.** Absolute concentrations of analytes from BMSC lysate treated with different conditions for 7 days.

| **Chemokine** | | | | |
|---|---|---|---|---|
| Marker | Group | Average Level (pg/µg) | SEM | P-value |
| IL-8 | BMSC | 0.001521 | 0.0005364 | 0.1097 |
| | BMSC+Treg | 0.006587 | 0.002413 | |
| IP-10 | BMSC | 0 | 0 | 0.1242 |
| | BMSC+Treg | 0.2084 | 0.1073 | |
| Eotaxin | BMSC | 0 | 0 | 0.3739 |
| | BMSC+Treg | 0.000168 | 0.000168 | |
| MCP-1 | BMSC | 0.07237 | 0 | 0.001 |
| | BMSC+Treg | 1.396 | 0.005664 | |
| MIP-1a | BMSC | 0 | 0 | 0.1025 |
| | BMSC+Treg | 0.01195 | 0.02404 | |
| MIP-1b | BMSC | 0 | 0 | 0.3739 |
| | BMSC+Treg | 0.01477 | 0.01477 | |
| RANTES | BMSC | 0 | 0 | 0.3739 |
| | BMSC+Treg | 0.3959 | 0.3959 | |
| **Growth Factor** | | | | |
| Marker | Group | Average Level (pg/µg) | SEM | P-value |
| bFGF | BMSC | 0.7561 | 0.05976 | 0.0575 |
| | BMSC+Treg | 7.806 | 2.668 | |
| G-CSF | BMSC | 0.006387 | 0.003247 | 0.0748 |
| | BMSC+Treg | 0.125 | 0.04941 | |
| GM-CSF | BMSC | 0 | 0 | 0.0939 |
| | BMSC+Treg | 0.0007847 | 0.0003586 | |

| | | | | |
|---|---|---|---|---|
| PDGF-bb | BMSC | 0 | 0 | >0.99 |
| | BMSC+Treg | 0 | 0 | |
| VEGF | BMSC | 0.08831 | 0.04508 | 0.0057 |
| | BMSC+Treg | 0.6617 | 0.09598 | |
| **Pro-inflammatory & Anti-inflammatory Marker** | | | | |
| Marker | Group | Average Level (pg/µg) | SEM | P-value |
| IL-1b | BMSC | 0.00231 | 0.0008215 | 0.0239 |
| | BMSC+Treg | 0.008411 | 0.001587 | |
| IL-2 | BMSC | 0 | 0 | 0.0074 |
| | BMSC+Treg | 0.00306 | 0.0006103 | |
| IL-6 | BMSC | 0.00811 | 0.001298 | 0.0107 |
| | BMSC+Treg | 0.6596 | 0.1444 | |
| IL-7 | BMSC | 0 | 0 | 0.3739 |
| | BMSC+Treg | 0.0004547 | 0.0004547 | |
| IL-9 | BMSC | 0 | 0 | >0.99 |
| | BMSC+Treg | 0 | 0 | |
| IL-12 (p70) | BMSC | 0.000652 | 0.000217 | 0.1362 |
| | BMSC+Treg | 0.003102 | 0.001299 | |
| IL-15 | BMSC | 0.04751 | 0.004995 | 0.0756 |
| | BMSC+Treg | 0.07894 | 0.0122 | |
| IL-17 | BMSC | 0.0001 | 0.0001 | 0.0287 |
| | BMSC+Treg | 0.004729 | 0.00138 | |
| IFN-ɣ | BMSC | 0.02365 | 0.001943 | 0.0088 |
| | BMSC+Treg | 0.1943 | 0.03563 | |
| TNF-α | BMSC | 0 | 0 | 0.0319 |
| | BMSC+Treg | 0.1236 | 0.03822 | |
| IL-1ra | BMSC | 0 | 0 | 0.0617 |
| | BMSC+Treg | 0.05079 | 0.01973 | |
| IL-4 | BMSC | 0.0003463 | 0.00009422 | 0.0137 |
| | BMSC+Treg | 0.002594 | 0.0005269 | |
| IL-5 | BMSC | 0.005119 | 0.003932 | 0.0178 |
| | BMSC+Treg | 0.102 | 0.02465 | |
| IL-10 | BMSC | 0 | 0 | 0.0917 |
| | BMSC+Treg | 0.02563 | 0.0116 | |
| IL-13 | BMSC | 0 | 0 | 0.0488 |
| | BMSC+Treg | 0.005193 | 0.001855 | |

**Table S3.** List of primers used for quantitative PCR analysis.

| Gene Symbol | Gene name | Assay ID |
|---|---|---|
| *GAPDH* | glyceraldehyde-3-phosphate dehydrogenase | Hs99999905-m1 |
| *RUNX2* | runt related transcription factor 2 | Hs01047973-m1 |
| *SP7* | Sp7 transcription factor | Hs01866874-s1 |
| *ALP* | alkaline phosphatase, liver/bone/kidney | Hs01029144-m1 |
| *BGALP* | bone gamma-carboxyglutamate protein | Hs00609452-g1 |
| *SPP1* | secreted phosphoprotein 1 | Hs00959010-m1 |
| *ITGB1* | integrin subunit beta 1 | Hs01127536_m1 |
